# Estimating Trends in Fertility in Kenya from Non Birth History Data

**DOI:** 10.1101/2020.09.09.289157

**Authors:** Paul Waweru Ngugi

## Abstract

This study aimed at determining the extent to which methods for estimating trends in fertility without use of birth history could be used on Kenyan surveys data by employing the own-children method (OCM) and reverse survival (RS) method in estimating fertility trend in the country. The study used data from 2015/16 Kenya Integrated Household and Budget Survey (KIHBS) and 2014 Kenya Demographic and Health Survey (KDHS). Data evaluation was done in order to obtain optimal fertility estimates. 2015/16 KIHBS data reported a Whipples index of 49.0 and 57.5 for terminal digits 0 and 5 respectively. Myer’s blended index was 2.9 and this was an indication that in general the data was accurate and therefore did not require any adjustment to improve its quality before use. Results from 2015/16 KIHBS showed that RS estimated Total Fertility Rate to be 3.5 as compared to OCM that estimated it to be 3.8. The results from 2014 KDHS dataset were consistent when using both RS and OCM. The two indirect methods can give consistent fertility estimates when the reference period is closer to the survey period but in the fourth and fifth year RS tends to systematically overstate fertility as compared to OCM. This study found out that in the absence of full birth history data, RS and OCM can reliably estimate consistent fertility estimates and trend.

## 1 Introduction

In a country, the composition, size and structure of a population depends on fertility, mortality and migration which are the components of population change. Fertility is known to have effects in the consumption of goods and services which include food, health, education, and housing. Malthus hypothesized that failure to control fertility would lead to high population growth and eventually reduce per capita income which might lead to a reduced consumption below subsistence level.

The Kenya government has continued to review and come up with better population policies, programmes and strategies aimed at addressing the challenges emanating from population management. For instance, in July 2012, the government adopted a population policy for national development. According to KenyaVision.2030 (2007), the aim of the policy was to ensure that the achievement of the economic, social and political pillars as depicted in Vision 2030’s are not hindered by population growth.

Casterline et al. (2017) observed that about four decades ago, Kenya had one of the highest fertility levels in the world, with Total Fertility Rate (TFR) being reported at 8.1 births per woman. Fig 1 shows that over the years there has been notable decline in fertility from 5.4 in 1993 to 4.6 births per woman in 2009. According to Kenya Demographic and Health Survey KDHS (2014), by 2014, TFR for the three years preceding the Kenya Demographic and Health Survey was 3.9 births for each woman, with rural women reporting at least one extra child compared to urban women.

**Fig. 1.**
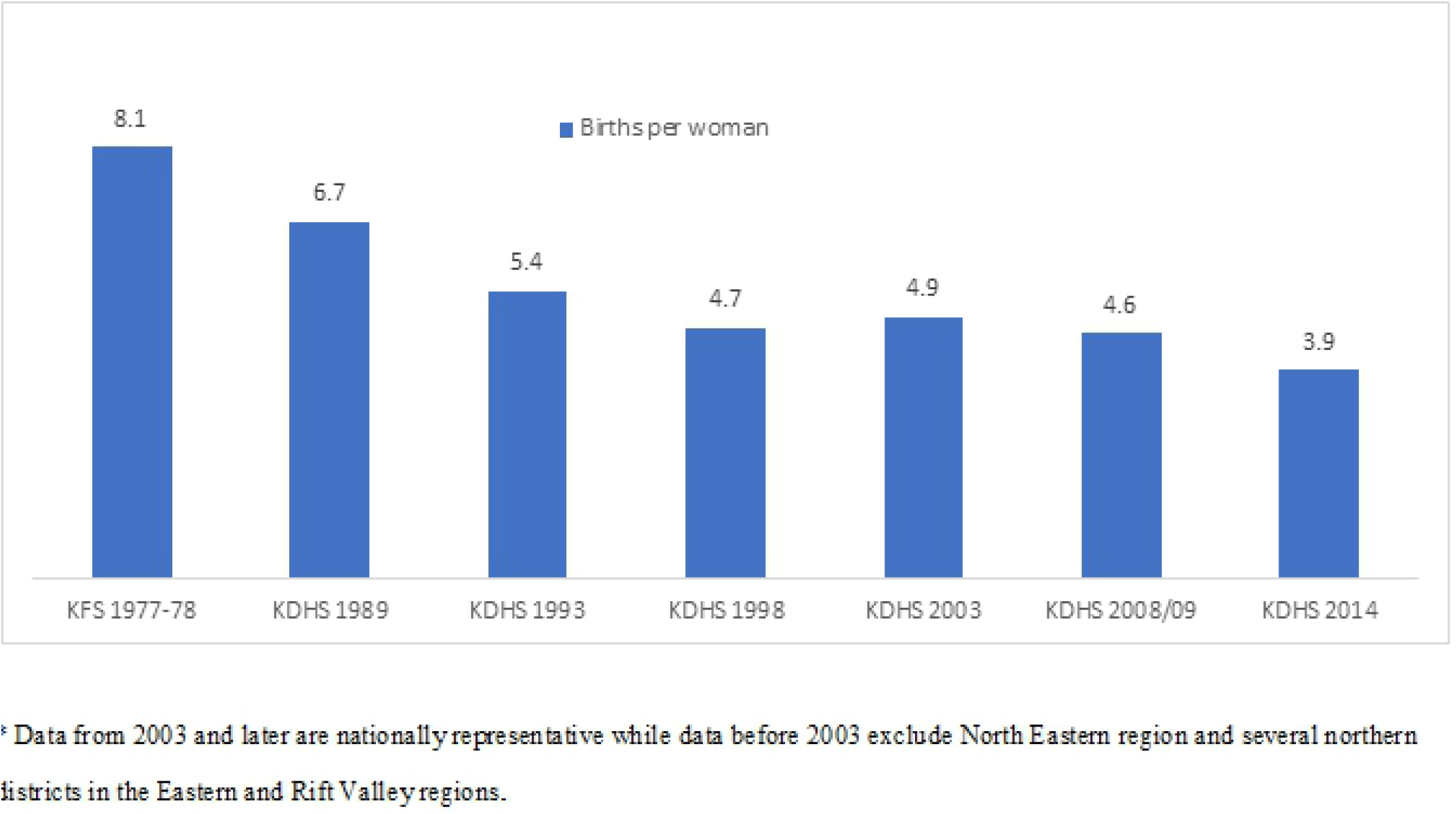
Total fertility rate trends, 1978-2014 Source:KNBS, 2015

Like other developing countries, Kenya does not have a vital registration system that is reliable which leaves surveys and censuses to provide fertility and child mortality estimates. In order to get estimates, various demographic techniques have to be applied. The type of data collected determines if direct or indirect techniques will be used to estimate measures of fertility and mortality. In Kenya, the main source of information on fertility rate is data about children ever born (CEB) alive and births that have occurred in a household in the 12 months preceding enumeration obtained from Demographic and Health Surveys (KDHS) and the country’s population census conducted every 10 years. The only available option of deducing past fertility trend from the available data is basically through comparison of the rates for the various survey/census years.

Besides KDHS, other surveys in Kenya do not employ birth history method when collecting data on fertility. Due to unavailability of such data, the OCM and RS provide an alternative indirect procedure of reconstructing the fertility experience of women up to a maximum period of 15 years preceding the enumeration. OCM makes use of information which is invariably obtained in a census or household survey with respect to age and sex of respondents, relationship among members of households and information on children who are enumerated with their mothers. Similarly, RS estimates fertility based on the population data collected from a census or a single-round survey as distributed by sex and age.

## 2 Own Children Method Application

When making inference for a population using information from the age distribution, OCM is the preferred technique since children are linked to their mothers using the mothers line number variable found in the household schedule (Cho et al., 1986). Through the use of this method, a mother’s and child’s age are used to determine the age of the woman during the time of giving birth to that child and also the time period probably when the same birth took place, hence providing a foundation for calculating age pattern fertility estimates; for children who may not be living with their biological mothers and in some cases where children could have died, then OCM advocates for making fertility level adjustments to the estimates.

Cho et al. (1986) noted that OCM was introduced to use population census data with the aim of generating differential fertility estimates. The method is applicable in any household survey where data on age and sex for everyone in the household has been collected. In additional OCM has the advantage of producing fertility estimates at single years and also for smaller population. OCM has been applied in past studies by Abbasi-Shavazi, (1997); Abbasi-Shavazi and McDonald (2002); Dubuc (2009); Coleman and Dubuc, (2010).

When studying differential fertility, Cho and Bogue (1970) marked that OCM has three advantages over other conventional methods. Firstly, OCM involves an additional step of coding or matching of biological mothers to their children. Secondly, the method allows the study of differential fertility for a period of 15 years preceding the census or survey period. Lastly, the method allows surveys to be modified hence making the data be available for studying fertility differentials. OCM was noted to be superior to data gotten from CEB as it can provide time associated fertility estimates. The technique can give current fertility rates as compared to CEB data that gives cumulative fertility. Cho and Bogue (1970) argued that over time cumulative fertility suffers from a lag in fertility change as compared to current fertility that gives results of actual changes in recent fertility.

Garenne and McCaa(2017) innovated an excel toolkit called 4 -parameter OCM which was an adaption of the original OCM and does not require use of external data sources when estimating adult mortality. Such appropriate innovations help in derivations of reliable fertility estimates that provide a basis of understanding of spatial variability of fertility regimes in an area and therefore is able to account for spatial disparities in adult mortality.

In the study of “Small Area Estimation of Fertility: Comparing the 4-Parameters OCM and the Poisson Regression-Based Person-Period Approach”, Ndagurwa and Odimegwu (2019) posited that 4-parameters OCM can obtain estimates of female adult mortality based on the survival status of children’s mothers which can be derived from a logit transformation of proportions who are maternal orphans,and by indirectly using the logits to estimate adult mortality. They noted that the viability in adult mortality for different geographic areas is reflected by the differences in the derived logits, slopes and intercepts.

OCM has the ability to estimate trends in fertility, Collins and Levin (2008) estimated TFR over time on four Kenyan censuses (1969, 1979, 1989 and 1999). Their results showed that TFR in Kenya increased from about 6 births per woman in 1950s and 1960s to about 8 births during the 1970s before declining to 6.5 births in the late 1980s. TFR was found to have reduced to about 5 births per woman in 1999. They noted that OCM had the strength of providing long term trends that act as external validity checks if more than one accurate data set is available. OCM is more reliable whenever fertility levels, trends and differentials are studied since it provides more revealing results as compared to conventional methods (Collins and Levin, 2008).

While carrying out an assessment of OCM of estimating fertility by birthplace in Australia, Abbasi-Shavazi (1997) pointed out that the technique had been applied to the 1986 data (Jain, 1989) and 1991 census (Dugbaza, 1994) with the aim of studying Aboriginal fertility. Abbasi-Shavazi made a conclusion that Jain and Dugbaza found out the OCM was able to produce reasonable fertility estimates of the Aboriginals.

Also, while examining data from the 2011 Iran census together with 2010 Iran DHS, Abbasi-Shavazi (2013), estimated fertility using OCM technique and compared the results with other direct and indirect methods of estimating fertility. The results showed that OCM was useful in estimating current differential fertility. He noted that other indirect methods rely on the assumption that fertility is constant for the period before enumeration. The assumption would mislead for countries still undergoing fertility transition.Their results showed that yearly fertility fluctuations could only be examined through the use of OCM.

## 3 Reverse Survival Application

When estimating for total fertility, RS model is preferred since its estimates are consistent and rarely suffer from any erroneous assumptions emanating from age distribution or if incorrect age patterns, mortality levels, trends are used during the estimation. When using RS, consistency of the estimates depends on the quality of data for both age and sex variables. When Spoorenberg (2014) was evaluating RS and how it estimates fertility, he noted that despite the technique being termed as simple and parsimonious to apply it is rarely used.

To show the contribution made by the method in estimating total fertility for past and present trends, Spoorenberg (2014) used data from five countries that included Kenya,Algeria,Ghana, Japan and Mongolia since these countries have distinct data quality issues and fertility levels. An excel template FE_reverse_4.xlsx, that is provided with Timæus and Moultrie (2012) was used to reverse survive a projected population that had been simulated by first projecting for more than 15 years through the use of sets mortality and fertility and also sex and age patterns.

Spoorenberg (2014) noted that RS gets estimates that are closer to original values gotten from the population. He concluded that sensitivity analysis done on the data showed that both incorrect age specific fertility patterns and mortality rates, trends and patterns do not affect RS estimates. RS suffers from effect of migration and therefore can only be applied at national levels unless there is detailed information about migration.

According to Timæus and Moultrie (2012), RS can derive fertility estimates just by the shape of the ASFRs without data on birth levels. The technique has the capability of stringing together reverse-survived estimates from successful surveys and census (Spoorenberg, 2014). Some of the limitations with RS are fertility estimates being biased by child under-reporting in general and mostly for children aged 0 and 1 years. To overcome the bias, fertility estimates can be aggregated by 5 year ages (United Nations, 1983). The other limitation is that the technique uses a constant age specific fertility rate though the shape of the ASFR curve is stable within a 10 year period. Lastly, RS technique does not estimate TFR by social characteristics, for example, education of mother.

## 4 Sources of Data

This study used data from the 2015/16 KIHBS and 2014 KDHS. KIHBS was an integrated household budget survey that was carried out in Kenya but was the first after the promulgation of the Constitution of Kenya which created devolved system of government. Both data sets were nationally representative surveys and were stratified based on the counties.

For KIHBS survey, women aged 12 years and above provided data on fertility and some of the eligible women were interviewed through proxies in case they were not present for interviews. In such cases information about the eligible women was provided by respondents who claim to have the best reproductive knowledge about them. According to Mislevy et al. (1991), accepting proxy answers is known to increase response errors in surveys.

On the other hand, KDHS collects both full birth history and summary birth history data. Therefore despite DHS data being expensive and time consuming to collect, it has the advantage of estimating fertility rates through direct and indirect techniques.

## 5 Data Quality

When carrying out research social scientists face many questions concerning the use of a particular data set, key among them is; does collected data suffer from errors and what could be the nature and magnitude, with specific reference being made to the main variables that matter most in the study? This question plays a vital role in this research, since the quality of collected data determines if the fertility estimates from such data will be useful and reliable.

This study used graphs, the age ratio method, Whipples and the Myer’s blended method and also the distribution of children ever born to evaluate age data. Data collected from surveys and census suffer from age errors as a result of coverage errors, which could emanate from failure by the enumerator to record the correct age as well as errors in age misreporting arising from the respondent. In particular, age data from sub saharan Africa and other developing countries is found to have high preference for some digits as a result of ignorance and illiteracy (Hlabana, 2010). In order to test age accuracy for data in single years, Myers’ blended index is mostly used because it aids in identifying if ages ending in some digits are preferred over others. The results from Myer’s index helps in identifying if the ten digits between 0 and 9 both being included are either preferred or avoided.

### Age Misstatements

Graphing of the population data as distributed by age in single years and by sex is beneficial to a researcher since any age heaping prevalent in the data is made visible at an early stage. Age heaping can be assessed visually and is deemed as a good indication of ages being piled together or heaped and is comparable to the assessment made from measures such as Whipple’s Index, Myers’ Blended Index and United Nations Age-Sex Accuracy Index. A manual from US Census Bureau concludes that while Whipple’s Index, Myers’ Blended Index and United Nations Age-Sex Accuracy Index are helpful as they provide summary statistics or useful during comparisons they fail to give an understanding into flaws that could be present in the data which are not obtainable through ratio and graphical analyses of the data. According to the US Census Bureau(1985) age heaping is exhibited by concentrating ages ending in 0 and 5 years for a given age distribution. Age heaping could equally occur for other ages based on how census data is collected or derived.

When demographic change is systematic due to absence of large exogenous events, the number of people found during the enumeration for all ages is expected to have a smooth progression. It is notable that fertility has always been high in developing countries and therefore population size is expected to decrease in a monotonic way by age. When number of births in absolute figures are found to have declined in recent years, then it is expected that children at younger ages would be slightly lower than at older ages.

During the 2015/16 KIHBS survey, women in the reproductive age groups were asked to report both their age with both month and year of birth questions being asked to act as a crosscheck. According to Shryock and Siegel (1976), the UN recommends that census and surveys should use either date of birth or age at last birthday when collecting data on age. It is noted that errors in age reporting are still persistent in both surveys and censuses.

According to Kpedekpo (1982) women in older ages are inclined to exaggerate their ages by overstating them. In many African societies, the elderly tend to be treated with high respect. This acts as a motivation for some older women to be inclined to exaggerate their age which comes with some associated respect. As a result, by exaggerating of the age of mothers this leads to some births being reported for women who are already above the reproductive ages which is mostly beyond 50 years and if fertility is estimated using conventional manner then they are consequently excluded (Zaba, 1981). In addition, this leads to fertility rates inflation reported by women beyond 40 years.

As earlier pointed out, graphing is another visual way to identify if the data has age misreporting. Fig. 2 shows population distribution in single years for both sexes. The graph shows that there was no strong preference for digit 0 and 5 as compared to the other digits as the determined peaks and troughs were also present in those other digits. This indicates that there was no significant age misreporting or heaping in the data.

**Fig. 2.**
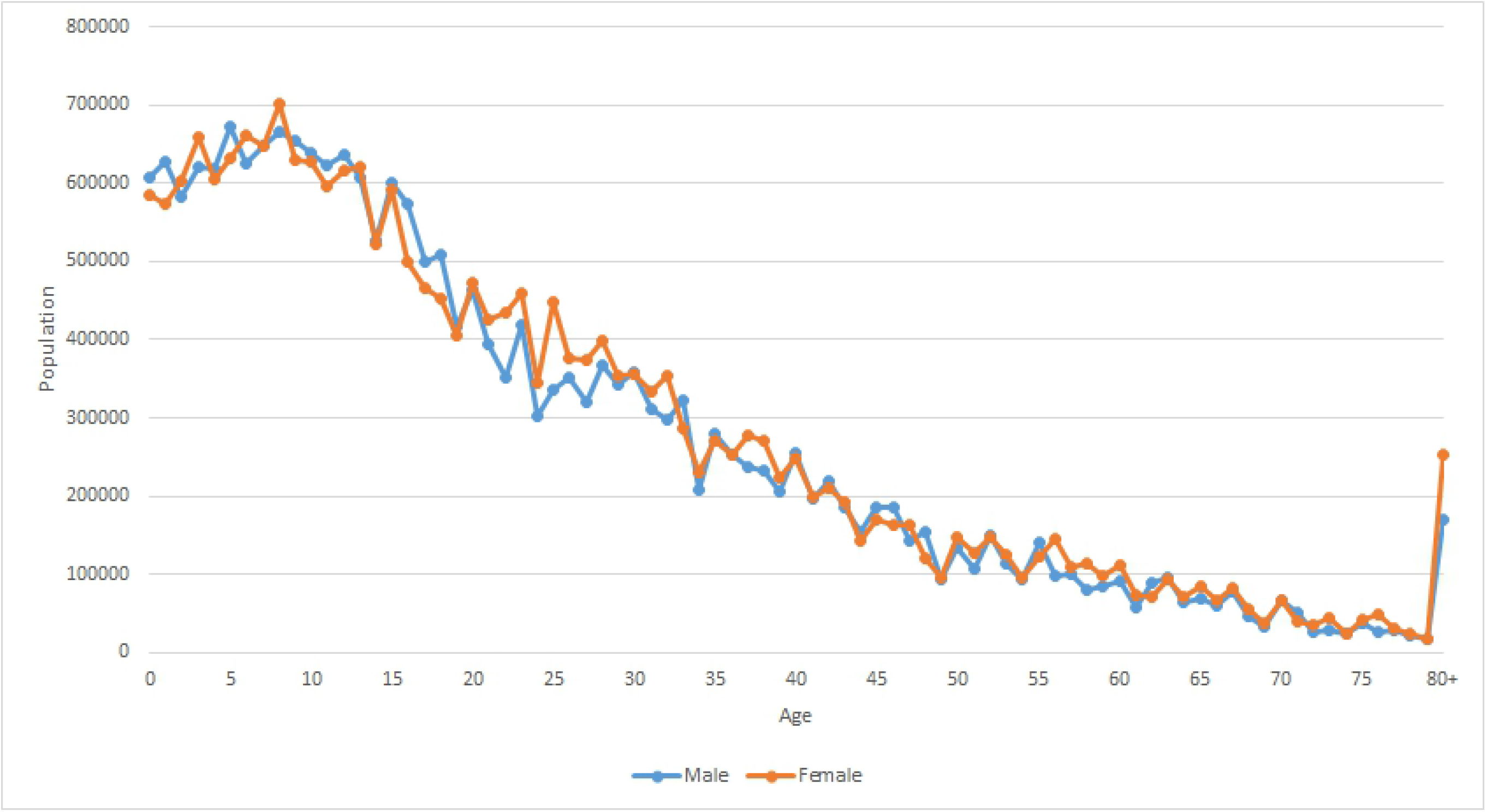
Age distribution in single years Source:Analysis of 2015/16 KIHBS

### Age ratios

Age heaping in some ages is easy to identify through graphs as compared to measures that need calculations. However, age ratio calculations tend to give an essential indication of probable undercounts or age displacements. Fig.3 indicates some minimal erratic overstating and understating of age for almost all the reproductive-aged women interviewed during 2015/16 KIHBS. This erratic behaviour for the younger and older women refutes the allegations that the data could have suffered from age understatement and overstatement. If age understatement or overstatement exists then it is expected to be seen over 100 for older and under 100 for younger women. The most noticeable age errors were seen for women in the 45-49 agegroup and 55-59 agegroup, with age reporting error being about 11%. The errors are assumed not to be systematic and therefore their effects on fertility estimates would cancel out.

**Fig. 3.**
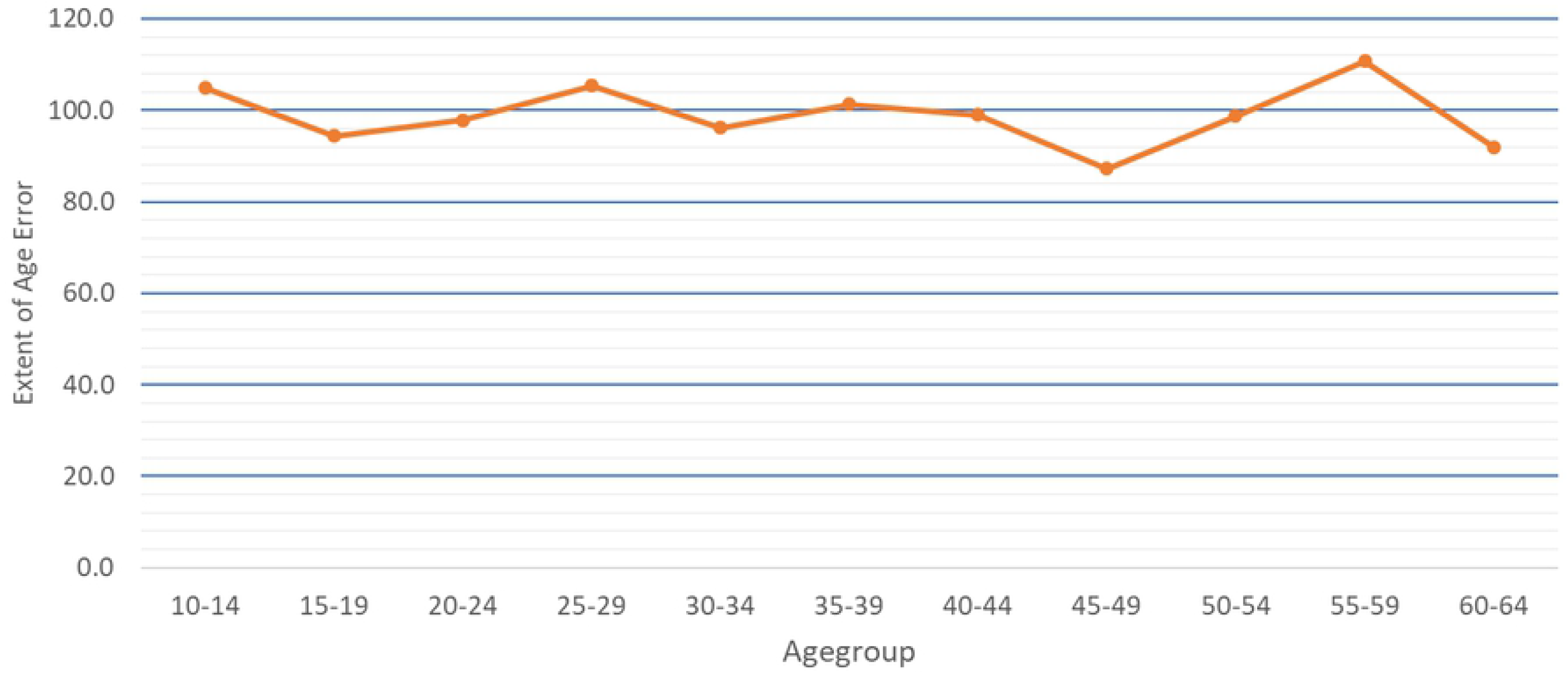
Extent of Age Error detection by agegroup Source:Analysis of 2015/16 KIHBS

### Age Heaping

Kpedekpo (1982) posits that some people have the tendency of showing preference in reporting certain ages over others. This tendency is called heaping or digit preference. While it is expected that digit preference can occur at any digit, but evidence shows that 0 and 5 digits are highly preferred. In order for avoidance or prevalence of each terminal digit to be detected, the Whipples index is employed. The value of the index calculated from the 2015/16 KIHBS data was 49.0 and 57.5 for terminal digit 0 and 5, respectively. This implies that the data was highly accurate and did not require adjustment before use.

However, Myer’s blended method is recommended in order to detect preference for all terminal digits. This method generates an index of preference for digit 0 to 9 which represents the deviation from 10% of the proportion of the total population that reports for each digit. From Table 1 and Fig.4 below, Myer’s blended method was calculated to be 2.9. The data shows insignificant preference and avoidance of ages for different terminal digits.

**Table 1.**
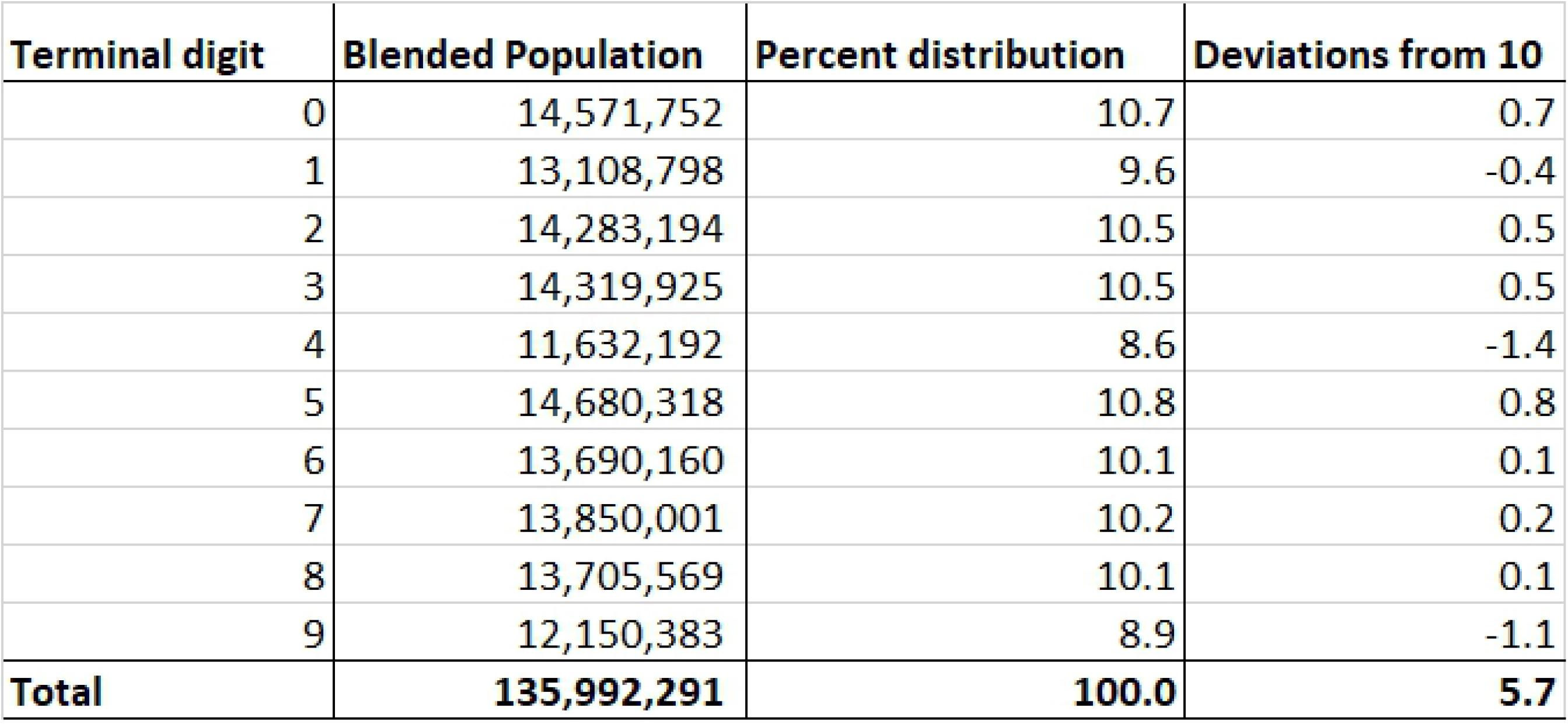
Results from Myer’s Blended Source:Analysis of 2015/16 KIHBS

**Fig. 4.**
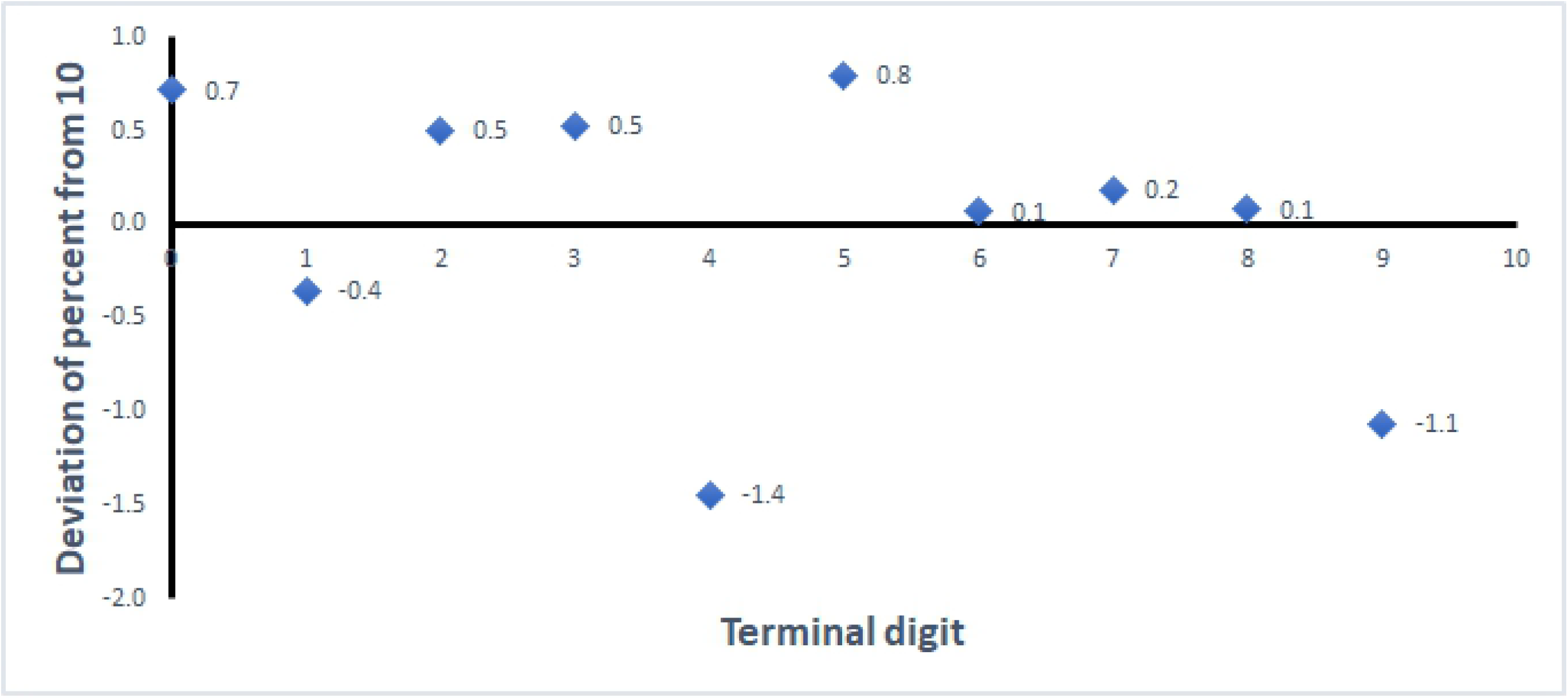
Myers index for digit preference,2015-2016 Source:Analysis of 2015/16 KIHBS

This study assumed that there was no significant missing data in the age variable and hence there was no imputation done in the analysis. Fig.5 depicts the distribution of Children Ever Born (CEB) by number of CEB. The figure shows that majority of the women reported to have ever given birth to 3 or less children, and the number of women decreases when the number of children increases.

**Fig. 5.**
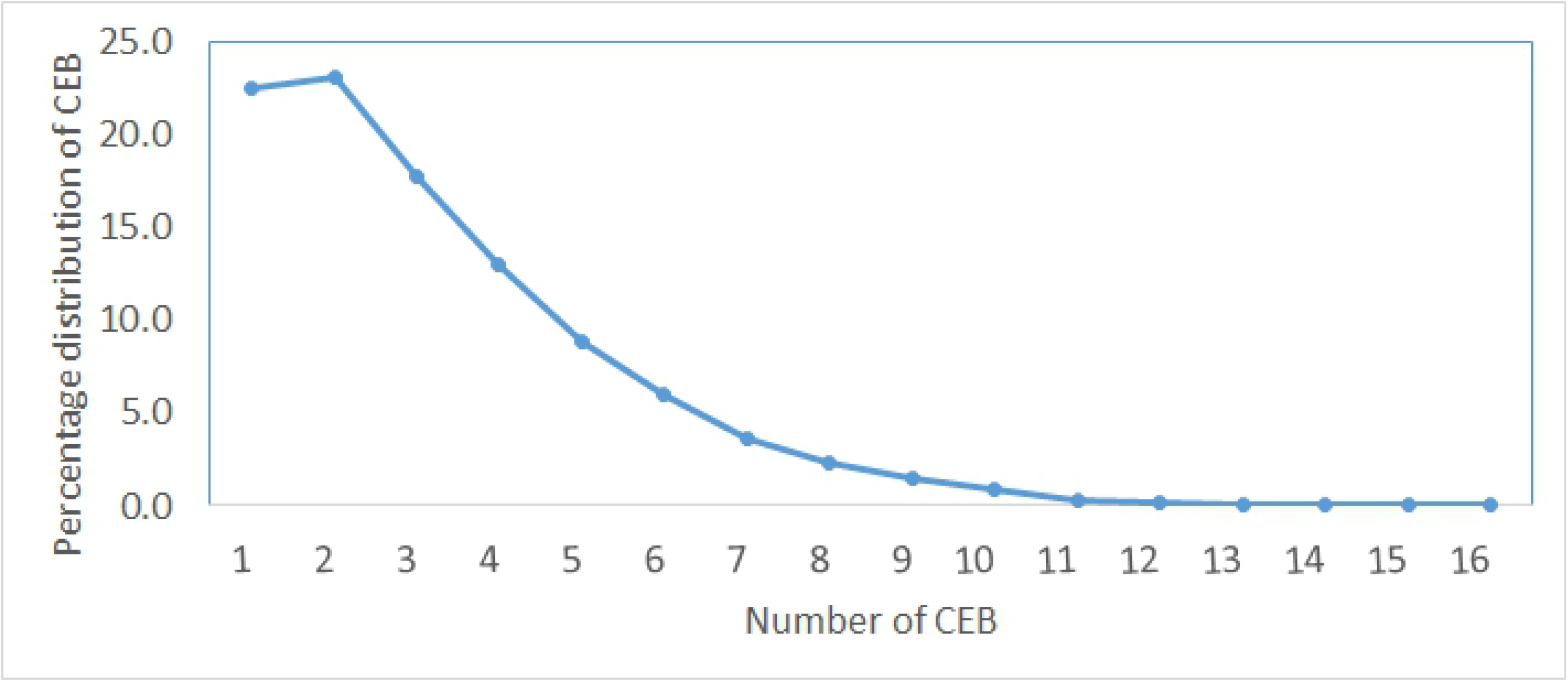
Distribution of Childen Ever Born Source:Analysis of 2015/16 KIHBS

In the 2015/16 KIHBS, 20,189 women in the 12-49 age bracket failed to report their parity. The failure to report parity might be as a result of errors during data entry for women with zero parity and is more prevalent to young women (United Nations, 1983). The age distribution of these women with zero parities did not show any bias and was represented across all the ages in the reproductive group. The proportion of women with zero parity was insignificant and were therefore not included in the analysis. From the analysis, parities varied from zero to 16 children. While it is expected that variation in fertility exists among different women, in 2015/16 KIHBS, some women reported CEB which was biologically impracticable as shown in Table 2 below. For example, there are 994 women aged 17 who reported to have had 12 children. There are other similar unrealistic cases that were reported and were therefore not used in the estimation of fertility since they would distort the estimates.

**Table 2.** Children Ever Born by Age of Mother Source:Analysis of 2015/16 KIHBS

| Age | Children born alive |  |  |  |  |  |  |  |  |  |  |  |  |  |  |  |  |
| --- | --- | --- | --- | --- | --- | --- | --- | --- | --- | --- | --- | --- | --- | --- | --- | --- | --- |
|  | 0 | 1 | 2 | 3 | 4 | 5 | 6 | 7 | 8 | 9 | 10 | 11 | 12 | 13 | 14 | 15 | 16 |
| 12 | 1,881 | - | - | - | - | - | - | - | - | - | - | - | - | - | - | - | - |
| 13 | - | 520 | - | - | - | - | - | - | - | - | - | - | - | - | - | - | - |
| 14 | - | 2,775 | 877 | - | - | 348 | - | - | - | - | - | - | - | - | - | - | - |
| 15 | 1,977 | 9,816 | - | - | - | - | - | - | - | - | - | - | - | - | - | - | - |
| 16 | 53 | 15,130 | 631 | 110 | - | 83 | - | - | - | - | - | - | - | - | - | - | - |
| 17 | - | 41,571 | 4,252 | 895 | 66 | - | - | - | - | - | - | - | 994 | - | - | - | - |
| 18 | 1,005 | 61,738 | 12,872 | 1,108 | 512 | - | - | - | - | - | - | - | - | - | - | - | - |
| 19 | - | 87,564 | 13,832 | 2,573 | 881 | - | - | - | - | - | 469 | - | - | - | - | - | - |
| 20 | - | 115,421 | 40,568 | 9,403 | 1,416 | 493 | 30 | 387 | - | - | - | - | - | - | - | - | - |
| 21 | - | 117,162 | 60,591 | 14,212 | 4,239 | 2,410 | 596 | - | - | - | - | - | - | - | - | - | - |
| 22 | - | 131,894 | 73,437 | 22,862 | 6,836 | 2,741 | - | - | - | - | - | - | - | - | - | - | - |
| 23 | 1,359 | 125,506 | 115,171 | 39,082 | 11,931 | 5,392 | - | - | - | - | - | - | - | - | - | - | - |
| 24 | 434 | 104,770 | 71,622 | 35,032 | 16,190 | 7,415 | 1,678 | - | - | - | - | - | - | - | - | - | - |
| 25 | 1,575 | 140,546 | 129,061 | 59,823 | 24,556 | 8,991 | 4,159 | 470 | - | - | - | - | - | - | - | - | - |
| 26 | 1,557 | 101,938 | 105,479 | 51,441 | 24,969 | 11,794 | 8,089 | 1,347 | 1,213 | - | - | - | - | - | - | - | - |
| 27 | 164 | 100,031 | 108,229 | 54,717 | 34,943 | 15,909 | 7,997 | 4,552 | 653 | 120 | - | - | - | - | - | - | - |
| 28 | 2,377 | 79,694 | 105,304 | 69,516 | 48,982 | 25,513 | 16,446 | 4,948 | 2,008 | 709 | - | - | - | - | - | - | - |
| 29 | 152 | 64,630 | 106,144 | 70,310 | 34,547 | 29,333 | 8,991 | 7,262 | 1,887 | 947 | - | - | - | - | - | - | - |
| 30 | 611 | 40,498 | 73,668 | 76,163 | 71,369 | 33,869 | 18,412 | 7,962 | 6,829 | 1,101 | 633 | - | - | - | - | - | - |
| 31 | - | 36,727 | 97,626 | 60,781 | 43,893 | 33,272 | 17,252 | 4,434 | 650 | 1,837 | 630 | - | - | - | - | - | - |
| 32 | 1,176 | 49,607 | 97,243 | 68,498 | 56,847 | 30,472 | 17,380 | 7,722 | 4,621 | 1,360 | 448 | 277 | - | - | - | - | - |
| 33 | - | 35,227 | 61,140 | 59,812 | 46,829 | 37,291 | 19,407 | 10,294 | 5,387 | 2,081 | 1,400 | - | 187 | - | - | - | - |
| 34 | 1,982 | 21,474 | 45,768 | 66,842 | 33,943 | 20,565 | 18,253 | 5,710 | 3,291 | 949 | 55 | 550 | - | - | - | - | - |
| 35 | 380 | 16,279 | 57,168 | 54,817 | 51,248 | 34,026 | 22,936 | 14,376 | 13,140 | 1,463 | 2,026 | 412 | - | - | - | - | - |
| 36 | - | 34,763 | 37,609 | 57,027 | 39,970 | 28,577 | 15,244 | 13,472 | 6,412 | 5,418 | 1,519 | - | 184 | - | - | - | 204 |
| 37 | - | 18,641 | 47,215 | 54,167 | 49,239 | 41,644 | 21,329 | 15,737 | 11,293 | 2,658 | 3,506 | 2,358 | 505 | - | - | - | - |
| 38 | - | 10,837 | 29,194 | 55,847 | 53,865 | 34,905 | 34,573 | 17,486 | 7,487 | 6,415 | 3,683 | 341 | 820 | - | - | 15 | - |
| 39 | - | 7,831 | 24,764 | 37,204 | 55,960 | 33,967 | 24,253 | 18,558 | 9,699 | 5,247 | 1,246 | 544 | - | - | - | - | - |
| 40 | 584 | 16,048 | 36,952 | 29,667 | 47,129 | 26,230 | 29,504 | 22,285 | 8,451 | 13,155 | 5,455 | 3,428 | 460 | 232 | - | - | - |
| 41 | - | 14,944 | 27,470 | 37,841 | 26,704 | 23,570 | 18,367 | 15,722 | 13,178 | 7,017 | 2,158 | 1,279 | 1,179 | - | 1,064 | 369 | - |
| 42 | 2,229 | 22,494 | 17,432 | 46,022 | 29,456 | 25,664 | 23,442 | 16,505 | 8,365 | 6,287 | 4,012 | 1,922 | 682 | 539 | - | - | - |
| 43 | 337 | 13,537 | 36,652 | 33,749 | 25,496 | 21,919 | 20,782 | 9,153 | 12,369 | 6,014 | 4,195 | 3,230 | 633 | - | - | - | - |
| 44 | - | 5,968 | 9,172 | 35,836 | 25,062 | 18,279 | 15,588 | 9,737 | 8,271 | 5,607 | 3,228 | 1,120 | 1,701 | 382 | - | - | - |
| 45 | 116 | 7,296 | 15,502 | 30,739 | 21,459 | 26,672 | 17,238 | 11,621 | 11,356 | 12,034 | 7,070 | 2,005 | 3,212 | 730 | - | - | - |
| 46 | 122 | 6,224 | 20,773 | 23,860 | 20,456 | 19,667 | 26,175 | 13,139 | 13,840 | 4,831 | 4,654 | 1,408 | 452 | 980 | 240 | - | - |
| 47 | 121 | 6,904 | 14,139 | 16,994 | 28,446 | 29,807 | 16,994 | 12,672 | 10,929 | 8,458 | 7,856 | 1,742 | 1,327 | - | - | - | - |
| 48 | - | 5,540 | 10,956 | 26,969 | 21,114 | 14,487 | 11,627 | 10,066 | 7,521 | 5,246 | 3,311 | 472 | 589 | 157 | - | - | - |
| 49 | - | 1,208 | 9,951 | 15,518 | 9,657 | 16,211 | 10,267 | 13,083 | 4,937 | 8,449 | 4,316 | 1,584 | 178 | - | - | - | - |

Fig.6 and Fig.7 show average parity as plotted by single ages and also by age groups, respectively. For both graphs, the average parity curves indicate a uniform increase in children ever born by age. Brass (1981), posits that average parity distribution should normally increase with age, and diminish towards the end of reproductive period. The average parity pattern was found to be acceptable for analysis. The average parities curve for CEB by single ages tends to meander from age 30 years as we tend to old ages. Unless when fertility is affected by irregular changes, the meandering indicates that as young women are consistent in the reporting of their parities as compared to older women. A curve depicting average parities by age-groups shows that there is uniform rate of increase in CEB as women move into higher age-groups. This figures show that when there are no changes in levels of fertility, average parities tend to increase with age since more women could be going back to childbearing. However the rate of increase tend to diminish as the women near the end of reproductive period since majority of them will be nearing the end of childbearing.

**Fig. 6.**
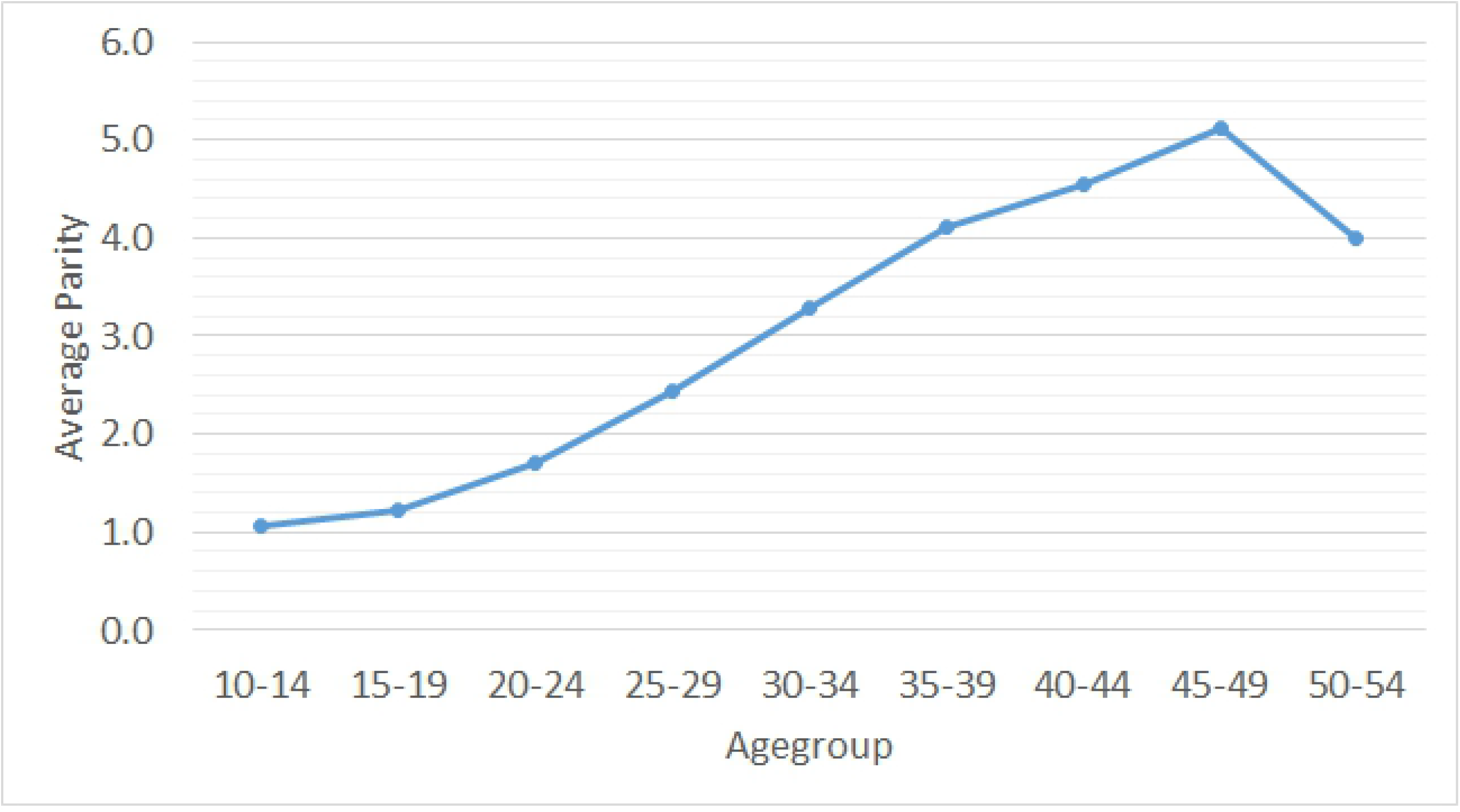
Average Parity distributions by Agegroup Source:Analysis of 2015/16 KIHBS

**Fig. 7.**
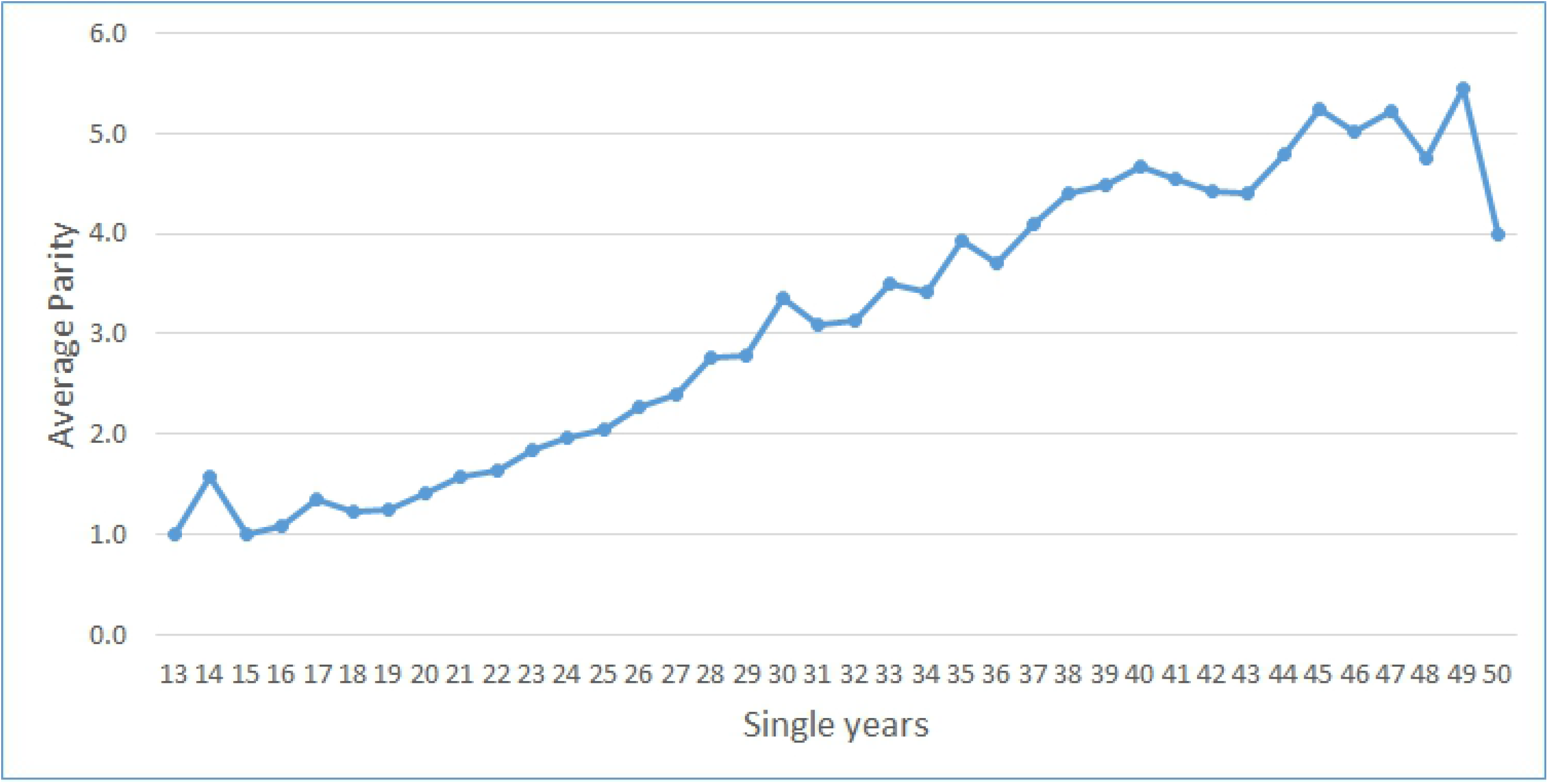
Average Parity Distribution by Single years Source:Analysis of 2015/16 KIHBS

2015/16 KIHBS shows that in total, 24,264,38612142 children were reported to have ever been born. However, when the women were asked detailed information on the number of children still living with them, number of children living elsewhere and and number of children who are dead by their sex orientation, a total of 24,339,839 children was reported as depicted in Table 3. The difference in reporting was 0.31 per cent makes the reporting of parities in the 2015/16 dataset satisfactory.

**Table 3.** Detailed Information on Total Number of Children Ever Born Source:Analysis of 2015/16 KIHBS

| **Variable** | **Cases** |
| --- | --- |
| Ever Given Birth | 7,464,876 |
| Never Given Birth | 5,313,596 |
| Children Ever Born | 24,264,386 |
| Sons Ever Born | 12,221,119 |
| Daughters Ever Born | 12,043,267 |
| Sons Living with Mothers | 9,824,729 |
| Sons Living Elsewhere | 1,746,142 |
| Sons Dead | 720,273 |
| Daughters Living with Mothers | 9,398,967 |
| Daughters Living Elsewhere | 2,009,160 |
| Daughters Dead | 640,569 |
| Total Children | 24,339,839 |

## 6 National estimates of total fertility rates (TFR) by OCM and RS

Fig.8 below shows the estimated levels and reconstructed trends of TFRs between 2001 and 2015. The results show that fertility trends using the RS techniques were systematically higher than those generated from the OCM models for years between 2001 and 2010. The two methods showed a uniform trends for the period between 2010 and 2015 but RS technique generated lower TFRs as compared to OCM. This shows that the two techniques are capable of producing consistent trends when the years are closer to the survey date as compared to when you move further from the survey date.

**Fig.8.**
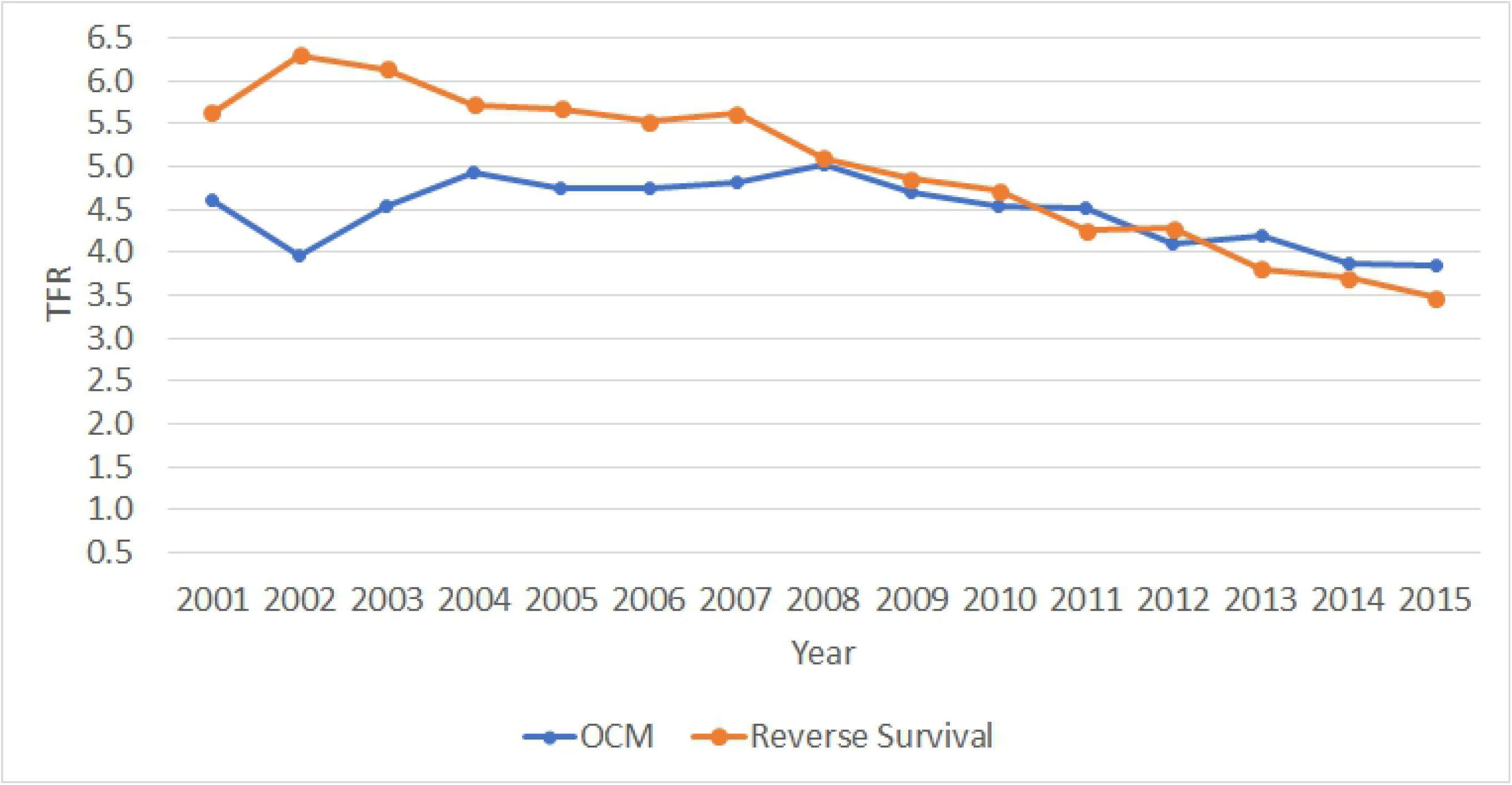
Reconstructed trends and estimates of TFRs Source:Analysis of 2015/16 KIHBS

Table 4 shows TFR and GFR results of the RS and OCM methods when applied to 2015/16 KIHBS dataset. The results show both techniques generated TFR estimates that ranged from 3.5 to 6.3 and 3.8 to 5.0 for RS and OCM, respectively, for the 2002 to 2016 period. In majority of the years, OCM had TFR estimates that were systematic lower than RS estimates. The fertility trend indicates that Kenya had a fertility stall in 2002 which supports the literature that Kenya experienced a fertility stall between 1993 to 2003 (Westoff et al., 2006) and (Mutuku, 2013). In the period under review, the average number of children being born to women aged 15 - 49 years was consistently low for OCM estimates.

**Table 4.**
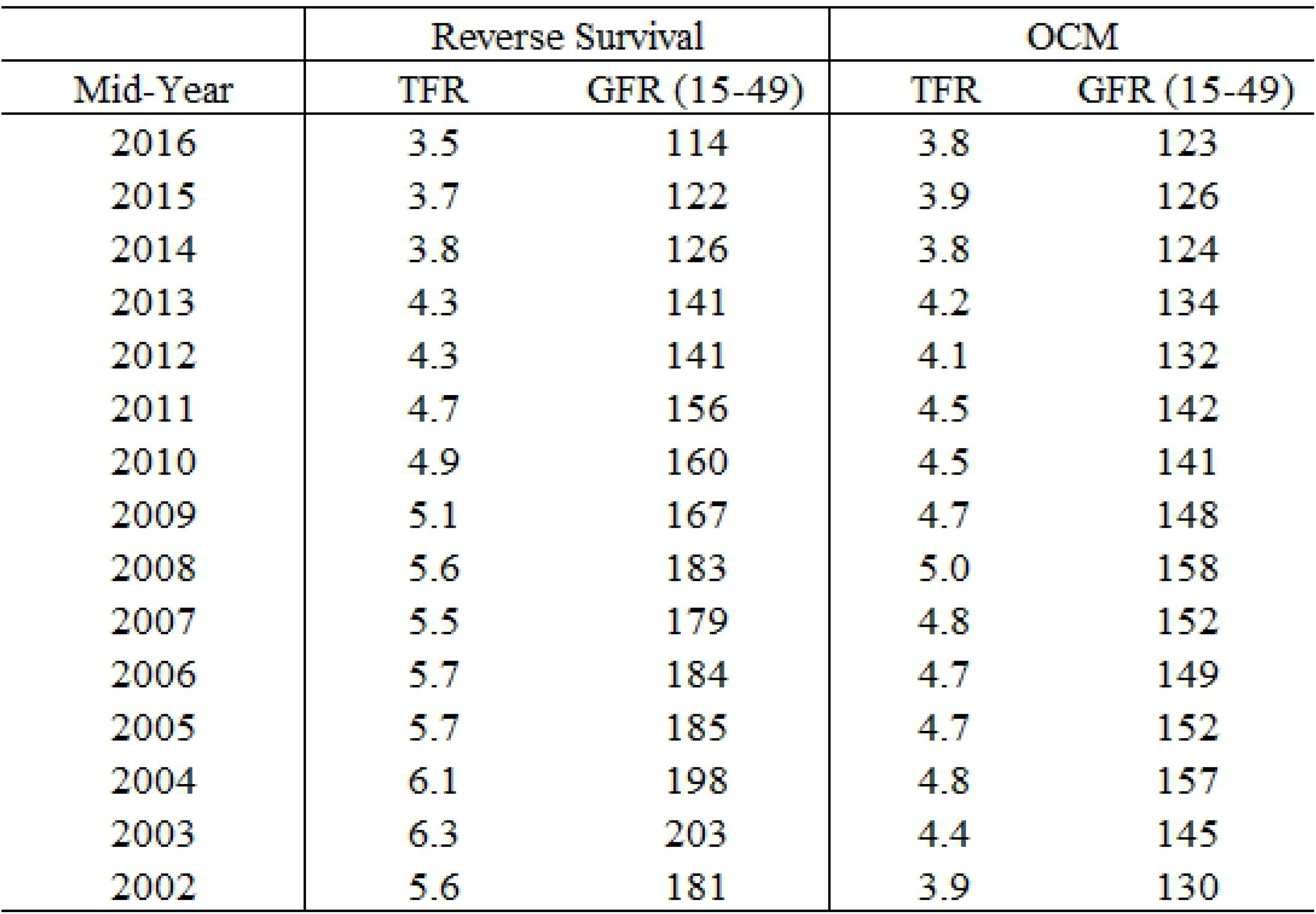
Fertility estimates per year(2002-2016) Source:Analysis of 2015/16 KIHBS

|  | Reverse Survival |  | OCM |  |
| --- | --- | --- | --- | --- |
| Mid-Year | TFR | GFR (15-49) | TFR | GFR (15-49) |
| 2016 | 3.5 | 114 | 3.8 | 123 |
| 2015 | 3.7 | 122 | 3.9 | 126 |
| 2014 | 3.8 | 126 | 3.8 | 124 |
| 2013 | 4.3 | 141 | 4.2 | 134 |
| 2012 | 4.3 | 141 | 4.1 | 132 |
| 2011 | 4.7 | 156 | 4.5 | 142 |
| 2010 | 4.9 | 160 | 4.5 | 141 |
| 2009 | 5.1 | 167 | 4.7 | 148 |
| 2008 | 5.6 | 183 | 5.0 | 158 |
| 2007 | 5.5 | 179 | 4.8 | 152 |
| 2006 | 5.7 | 184 | 4.7 | 149 |
| 2005 | 5.7 | 185 | 4.7 | 152 |
| 2004 | 6.1 | 198 | 4.8 | 157 |
| 2003 | 6.3 | 203 | 4.4 | 145 |
| 2002 | 5.6 | 181 | 3.9 | 130 |

Fig. 9 presents a comparison for 5 year averages of ASFRs using OCM technique, for both 2015/16 KIHBS and 2014 KDHS. The ASFRs obtained from the 2015/16 KIHBS and 2014 KDHS show a uniform trend for ages 12 to 49 years. KIHBS data produced ASFRs that were lower for women aged 15 to around 25 years. The two dataset reported similar ASFRs for the women aged betwen 25 and 35 years. Fig.9 also shows that the trends changes for women aged 35 to 49 years where 2014 KDHS dataset reports slightly lower ASFRS. The figure shows that OCM produces schedules of ASFRs that are stable based on the smoother graphs. The ability of OCM to produce smoother schedules of ASFRs gives it some strength that can be exploited at lower levels of aggregation to produce localised fertility estimates. The consistent schedules of ASFRs from Fig.9 imply that fertility levels in Kenya obtained from 2015/16 dataset are consistent and therefore reliable.

**Fig. 9.**
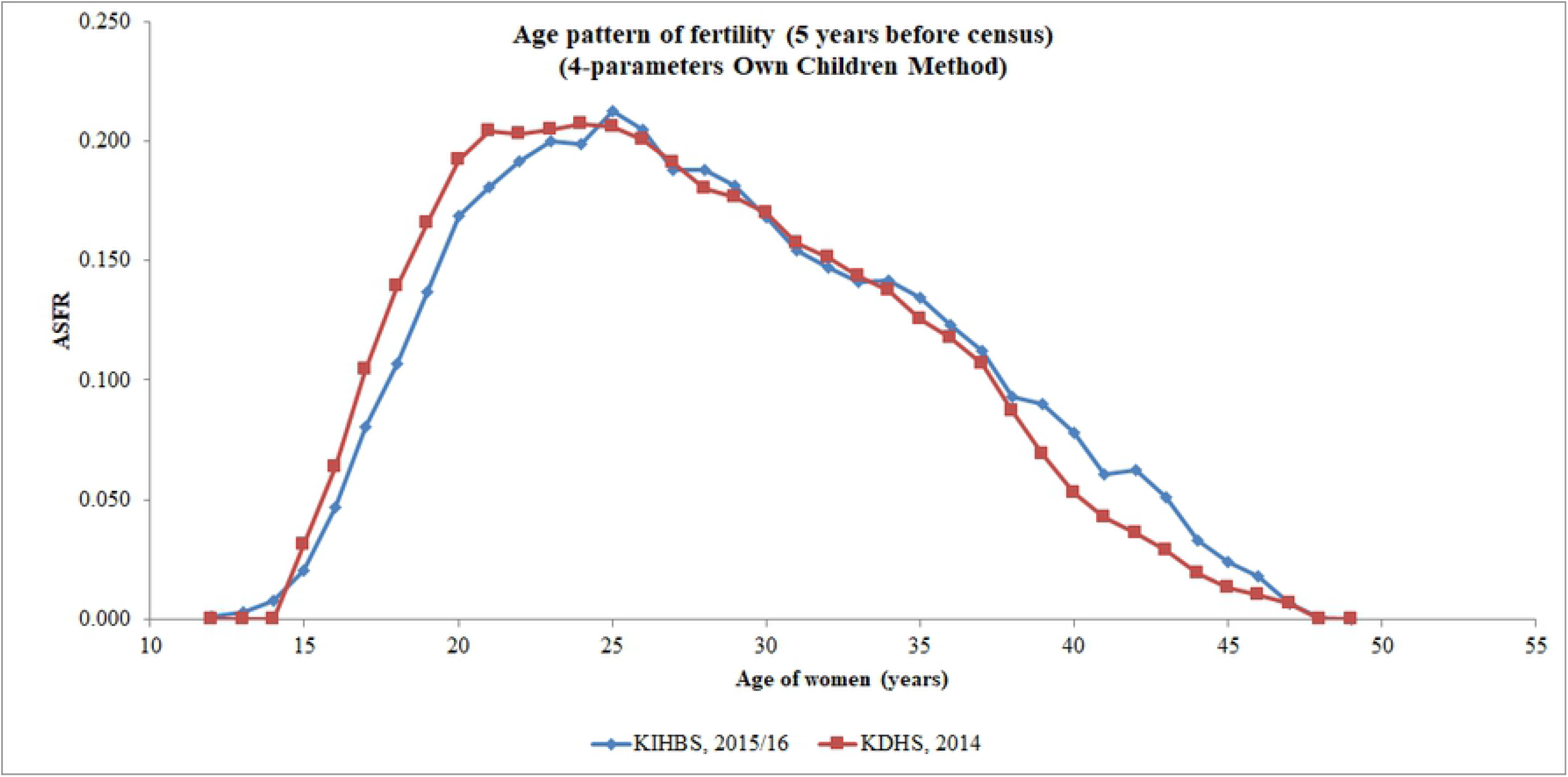
Comparison for 5 year Averages of ASFRs using OCM technique Source:Analysis of 2015/16 KIHBS

The comparison of TFRs obtained from both 2015/16 KIHBS and 2014 KDHS datasets using OCM and RS is presented in Fig. 10. The estimated levels and reconstructed trends of TFRs indicate that the two methods produces a similar and uniform trend for the five years preceding both surveys period. For the remaining TFRs estimated using the RS were consistently higher than those obtained from the OCM technique which is expected. Fig. 10 shows that in the fourth and fifth years preceding the survey, the gap between TFR estimates from the two methods appears to widen.

**Fig. 10.**
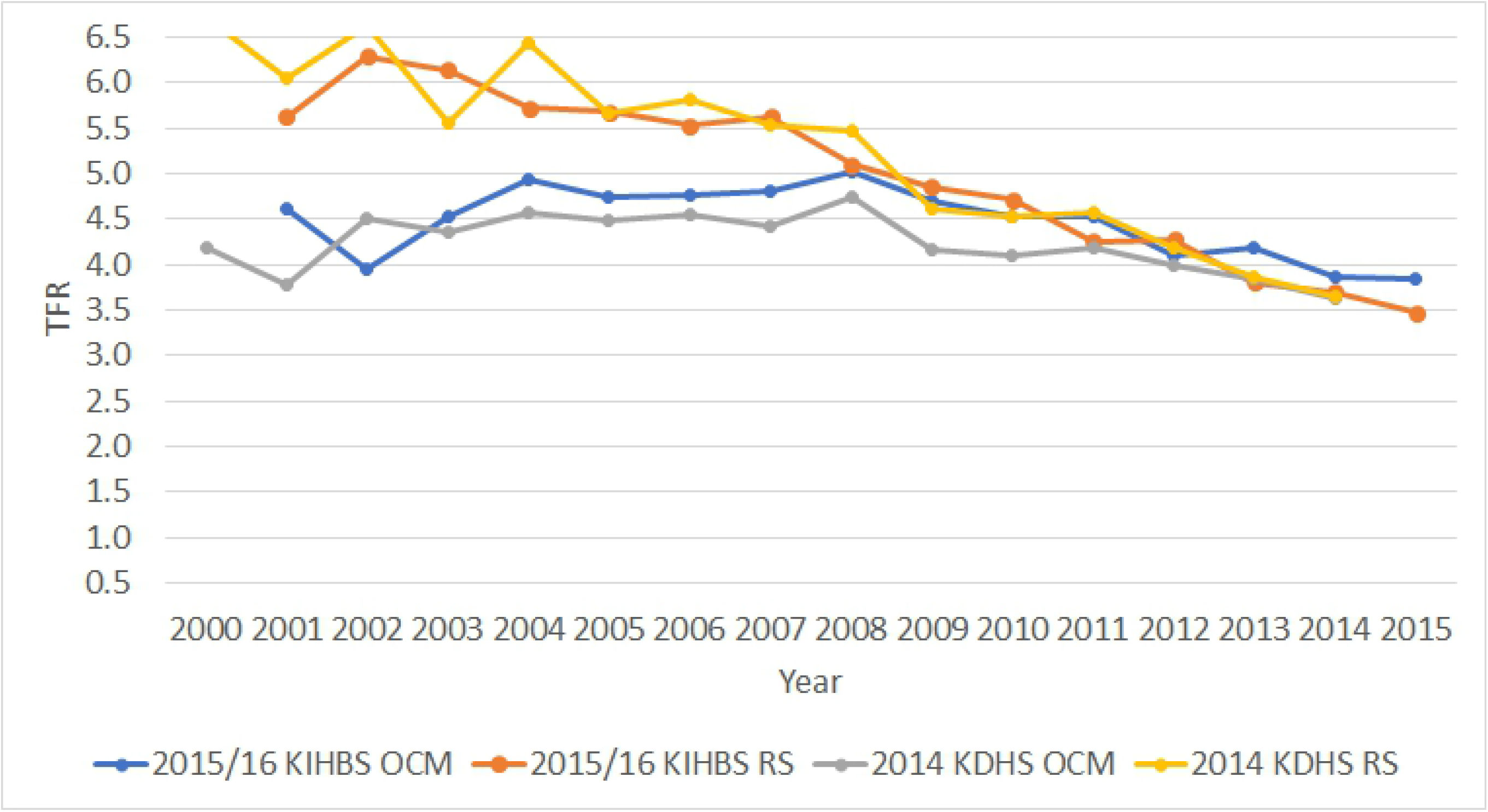
Comparison TFRs obtained from 2015/16 KIHBS and 2014 KDHS Source:Analysis of 2015/16 KIHBS and 2014 KDHS

Fig. 11 depicts a comparison of TFRs obtained from 5 year groups using OCM for both 2015/16 KIHBS and 2014 KDHS dataset. The TFRs obtained display a uniform trend for all the years preceding survey years, that is from 2001 to 2013. This indicates that the results obtained from OCM are consistent despite using different datasets which gives an indication that the estimated TFRs are reliable.

**Fig. 11.**
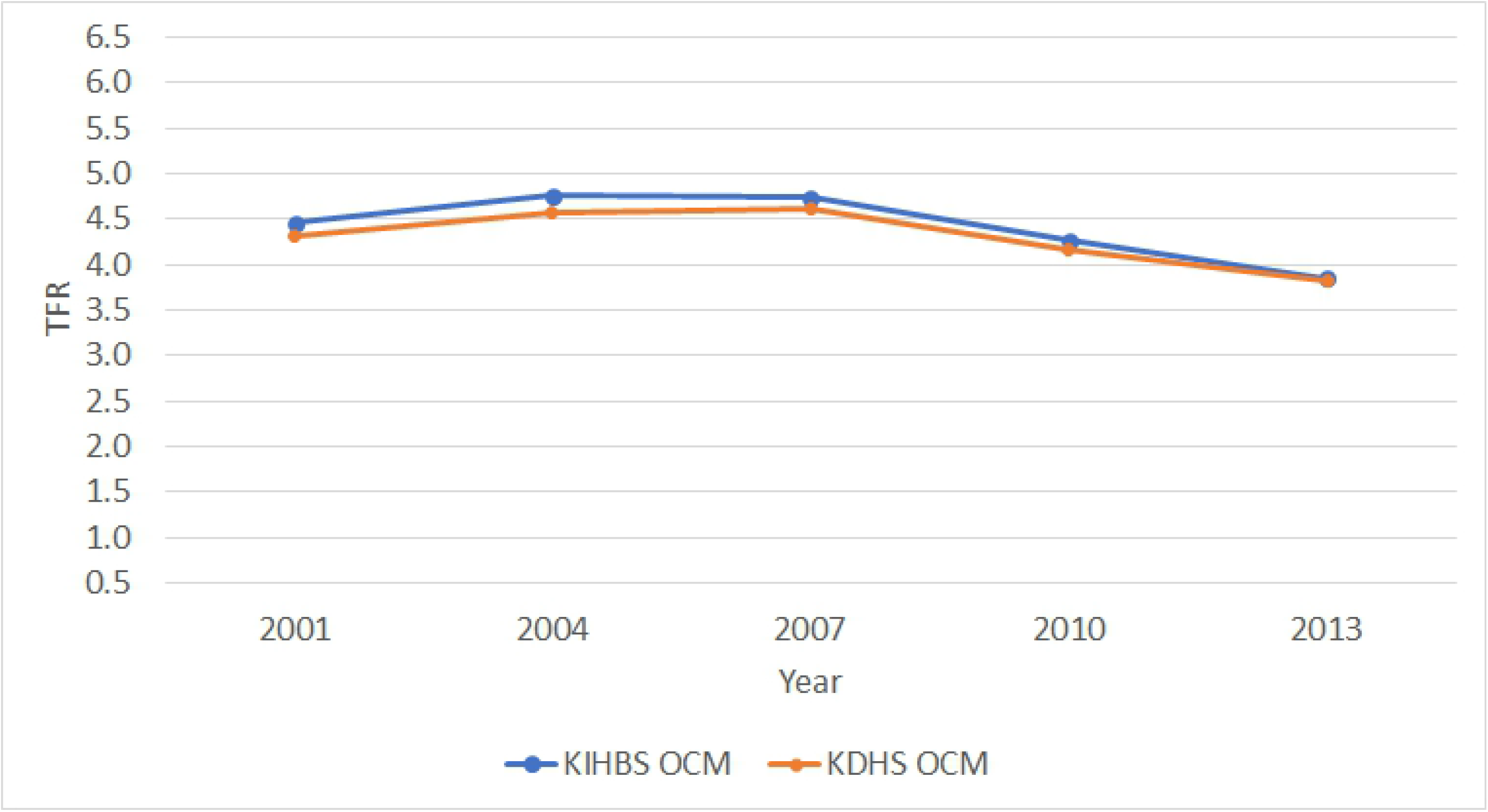
5 Year groups OCM Source:Analysis of 2015/16 KIHBS and 2014 KDHS

Table 5 shows the estimated results of the RS and OCM method when applied to 2014 KDHS dataset. The results indicate that TFR estimates range from 3.6 to 6.7. Both RS and OCM reported TFR of 3.6 in 2014 as compared to birth history method that was reported at 3.9. The results supported what Avery et al. (2013) had posited that FHB produces significantly larger TFR as compared to OCM technique. The results also showed that as you move away from the survey period, RS produces reconstructed TFR estimates that are systematically higher than OCM. The estimated TFR from 2014 KDHS were similar to those from 2015/16 KIHBS indicating that the estimates from the latter are reliable.

**Table 5.**
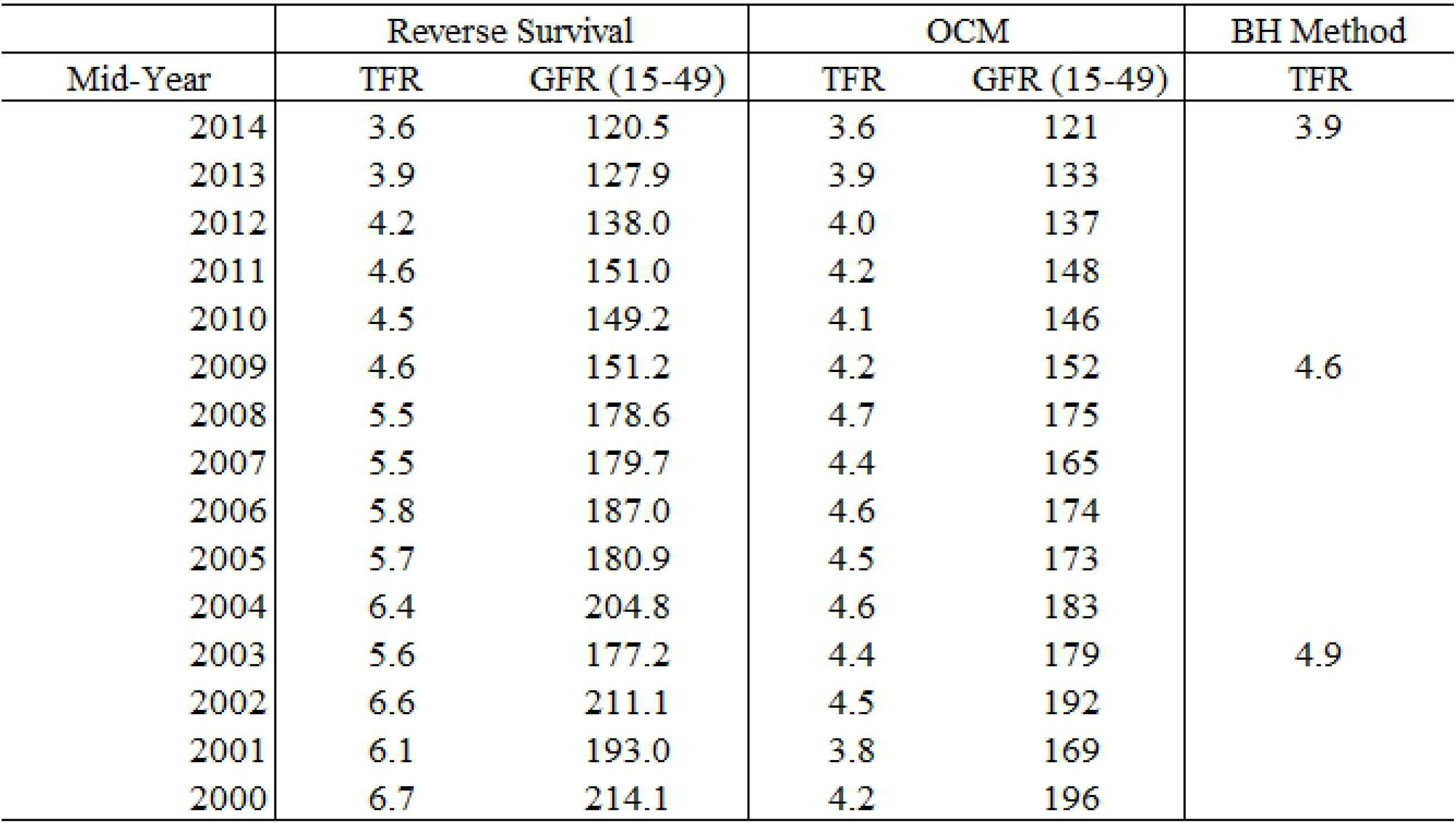
Fertility estimates per year(2000-2014) Source:Analysis of 2014 KDHS

## 7 Discussion

The purpose of this study was to determine the extent to which methods for estimating trends in fertility without use of birth history could be used on Kenyan surveys data. Results from 2015/16 KIHBS showed that RS estimated TFR to be 3.5 as compared to OCM that estimated it to be 3.8. To check for the consistency of the results, 2014 KDHS was used as shown in Table 5. The results from 2014 KDHS dataset are consistent when using both RS and OCM but under-reports TFR compared to birth history method. Both Table 4 and 5 indicate that even though the two models have different assumptions, derived estimates are within range.

The study showed that both techniques can reveal a trend in fertility for upto a period of 15 years. The results show that RS can produce better current estimates as compared to OCM though both methods have an added advantage since they can reveal a trend in fertility.

The results from this study agree with what Spoorenberg (2014) had noted that RS gets estimates that are robust and closer to what other indirect methods get. The results also confirm the notion that OCM has advantages over other indirect methods due to its non requirement of prior assumptions on past vital rates and also does not require any direct and detailed information on birth statistics. This study was an extension of the study done by Collins and Levin (2008). They had used the original OCM, while this study was based on the 4 parameter model developed by Garenne and McCaa(2017).

## 8 Conclusion

Based on the results, this study concluded that both RS and OCM can use non birth history data and reliably estimate trends in fertility levels in Kenya. The two indirect methods can give consistent fertility estimates when the reference period is closer to the survey period but in the fourth and fifth year RS tends to systematically overstate fertility as compared to OCM and therefore the latter is preferred since it can give reliable and consistent current estimates and trends. Since estimates based on such methodologies compare with those from the full birth history method, it can be concluded that the non birth history data can be used to estimate current fertility and also produce a fertility trend for upto 15 years. When the two techniques were applied to the 2014 KDHS dataset they were found to under-report TFR compared to birth history method.

## Bibliography

Abbasi-Shavazi, M. (1997). An assessment of the own–children method of estimating fertility by birthplace in australia. Journal of the Australian Population Association, Vol. 14, No. 2 (November 1997), pp. 167–185.

Abbasi-Shavazi, P McDonald, M. H.-C. (2013). Assessment of the own-children estimates of fertility applied to the 2011 iran census and the 2010 iran-midhs. In XXVII IUSSP International Population Conference, Busan, Korea, 26-31 August 2013.

Abbasi-Shavazi, M. J. and McDonald, P. (2002). A comparison of fertility patterns of european immigrants in australia with those in the countries of origin. Genus, 58(1):53–76.

Avery, C., Clair, T., Levin, M., and Hill, K. (2013). The ‘own children’ fertility estimation procedure: A reappraisal. Population Studies, 67(2), 171–183. https://doi.org/10.1080/00324728.2013.769616.

Casterline, J. B., Bongaarts, J., Askew, I., Maggwa, N., and Obare, F. (2017). Fertility Transitions in Ghana and Kenya: Trends, Determinants, and Implications for Policy and Programs. Population and Development Review, 43:289–307.

Cho, L., Retherford, R., and Choe, M. (1986). The own-children method of fertility estimation. www.popline.org, Honolulu, Hawaii, The East-West Center, 1986. xvi, 188p.

Cho, L. J., W. H. G. and Bogue, D. J. (1970). Differential current fertility in the united states. Chicago: University of Chicago.

Collins, O. and Levin, M. J. (2008). Fertility levels, trends and differentials in kenya: How does the own-children method add to our knowledge of the transition? African Population Studies.

Dubuc, S. (2009). Application of the own-children method for estimating fertility by ethnic and religious groups in the uk. Journal of Population Research, 26(3):207–225.

Dugbaza, T.. A. B. o. S. D. S. (1994). Recent trends and differentials in aboriginal and torres strait islander fertility, 1986-1991. Canberra : Australian Bureau of Statistics, Demography Section, 1994.

Hlabana, T. (2010). Application of the p / f ratio method in estimating fertility levels in lesotho.

Jain, S. (1989). Estimation of aboriginal fertility, 1971-86: An application of the own children method of fertility estimation. Canberra : Australian Bureau of Statistics.

KDHS (2014). Kenya Demographic and Health Survey 2014. Technical report, Rockville, MD, USA.

KenyaVision.2030 (2007). Kenya vision 2030. Government of the Republic of Kenya, Ministry of Planning and National Development and the National Economic and Social Council (NESC), Office of the President, Nairobi.

Kpedekpo, G. M. K. (1982). Essentials of demographic analysis for Africa.

Mislevy, R. J., Rubin, D. B., and Little, R. J. A. (1991). Statistical analysis with missing data. Journal of Educational Statistics, 16(2):150–155.

Ndagurwa, P. and Odimegwu, C. (2019). Small area estimation of fertility: Comparing the 4-parameters own-children method and the poisson regression-based person-period approach. Spatial Demography.

Shryock and Siegel (1976). The methods and materials of demography. Academic Press, San Diego.

Spoorenberg, T. (2014). Reverse survival method of fertility estimation: An evaluation. Demographic Research, 31:217–246.

Timæus, I. and Moultrie, T. (2012). Estimation of fertility by reverse survival. Tools for Demographic Estimation. Paris: International Union for the Scientific Study of Population.

United Nations (1983). Manual ten. Population studies; no. 81.

